# High resolution single-cell chromatin 3D modeling reveals coherent chromatin aggregation with varied structures in controlling genome function stability

**DOI:** 10.1101/546572

## Abstract

The genome 3D architecture is thought to be related to regulating gene expression levels in cells and can be explained by genome-wide chromatin interactions which have been explored by chromosome conformation capture based techniques, especially Hi-C. Based on single-cell Hi-C data, we developed a new method in constructing experimental consistent 3D intact genome structures for individual cells with a resolution of 10kb or higher. The modeled structures showed marked variations of 3D genome organization across different cells. However, chromosome loci marked by different proteins, such as CTCF and post-translationally modified histones, are consistently non-specifically aggregated in space. Interestingly, similar aggregations between active enhancers and active promoters were observed, especially for those separated by genomic regions of the scale of megabase or larger. Such long-range associations between active enhancers and promoters are strongly correlated with spatial aggregation of chromosome loci marked by different proteins. Through analyzing the 3D structures of intact genome, we proposed that coherent gene activation profiles among individual cells can be achieved by the consistent aggregation of protein marked loci instead of maintaining identical folded conformations.

---

Genome architecture that are characterized by interactions between chromosome regions separated by DNA sequence of certain length is believed to play important roles in controlling and regulating biological functions in cell nucleus^1–5^. Hi-C experiments based on genome-wide DNA fragment digestion and ligation make it possible to capture these interactions and construct contact frequencies by examining DNA sequences organized in close spatial proximity^6–11^. The contact frequency matrix generated by Hi-C experiment shows hierarchical architecture of genome from conserved topological associating domains (TAD) at hundreds of kilobase scale^12–15^ to spatially segregated A/B compartments at the megabase scale^6,8,12,16,17^. Alternation of this architecture, such as deletion of TAD boundaries, can lead to dysregulation of gene expressions in cells^14,18^. Starting from Hi-C contact matrix, many efforts have been made in developing computational methods that are able to construct models consistent to Hi-C experiments^6,19–28^. In these methods, single chromosome is coarse-grained as a polymer chain and ensembles of chromosome 3D structures are generated after applying physical interactions between monomers along the chain. These modeled structures of these methods greatly enhance our understanding of chromosome folding and reveal novel principles of structural organization in regulating biological functions of chromosomes. However, recent studies have shown that genomes in single-cell level are organized in varied structures, which challenges the principles revealed by populated Hi-C based models^21,29^. To investigate genome folding in individual cells, Stevens *et al*. designed new procedures of chromosome conformation capture experiments in single-cell level and structures were calculated to provide insight into 3D organization of intact genome in single nucleus for the first time^29^. Nonetheless, due to the relative low resolution (100kb), these structures provide limited information of interactions among chromosome loci that is only available at higher resolution; these information is critical in studying the regulation of genome folding on gene expression. Therefore, we developed a new method aimed at obtaining high-resolution structures of intact genome from data obtained in single-cell Hi-C experiment. The calculated structures not only exhibit consistent features compared to both theoretical and experimental observations, but also provide evidence for principles of genome folding via non-specific aggregation of chromosome loci marked by different proteins, as well as promoting the facts that such aggregations is important in the association between active enhancers and promoters at genome-wide scale. Based on our modeled structures, we therefore propose a new mechanism of genome folding that integrate varied folded structures and consistent biological functions of individual cells.

## Result

### Construction of high-resolution genome architecture from single-cell Hi-C

We developed a new method to construct three-dimensional structure of intact genome using genome-wide contact data generated by Stevens’ single-cell Hi-C experiments ^29^. In our method, each chromosome was presented by a polymer chain with each monomer representing one chromosome locus along the sequence. By maximizing the coherence between polymer chain conformations and single-cell Hi-C data, we constructed high-resolution structures of intact genome for 8 individual mouse embryonic stem cells.

Convergent 3D structures were constructed at the genomic resolution of 10 kb for different cells (ensemble averaged rmsd of 10.5 particle radii for cell 1). These 10kb binned structures and the recalculated 100-kb structures both exhibited similar spatial organization when comparing to those in Stevens’ work^29^(shown in Fig.1a, averaged rmsd < 6 particle radii for 100kb resolution), which validated the reliability of both calculated structures and the strategy in constructing these structures. The contact matrices reconstructed based on modeled structures successfully reproduced the interaction patterns between genome-wide loci presented in Hi-C data (>99.8% Hi-C contact pairs reproduced, Fig.1b, Extended Data Fig.1a) and the violated loci pairs were more frequently observed to locate at sequentially close regions (Extended Data Fig.1d). Besides, loci pairs of 20 times more population were identified in modeled structures when comparing to experimental detection, suggesting the number of detected contact reads obtained by current experiment technique is far from upper limit of the possible read number in one cell nucleus. More strikingly, recalculation after removing 90% Hi-C data (with contact pair number < 10,000 after removal) still generated structures with similar contact patterns (Fig.1a, Fig.1b), suggesting the high efficiency and robustness of our method in modeling genome structures. The calculation for recombined Hi-C data of two different cells leaded to highly stretched structures with a 20-fold increase in the number of violated contact loci pairs (11.6% pairs violating experimental derived constraints, Extended Data Fig.1d) and larger degrees in violating the distance restraints presented in merged contact matrix (maximum distance of 114 particle radii for violated pairs, compared to 7.7 particle radii for separated data). In addition, we constructed 1kb-resolution conformations for each cell satisfying constraints of Hi-C contacts. Note that these high-resolution structures were not used in further analysis due to the relative large conformational variation among replicated structures derived from limited amount of contact reads (rmsd of 13.3 particle radii). Taking together, the high consistency between calculated structures and experimental data strongly validated the reliability and robustness of our method in utilizing single-cell Hi-C data to reproduce the folded structure of intact genome at high resolution level.

**Figure 1.**
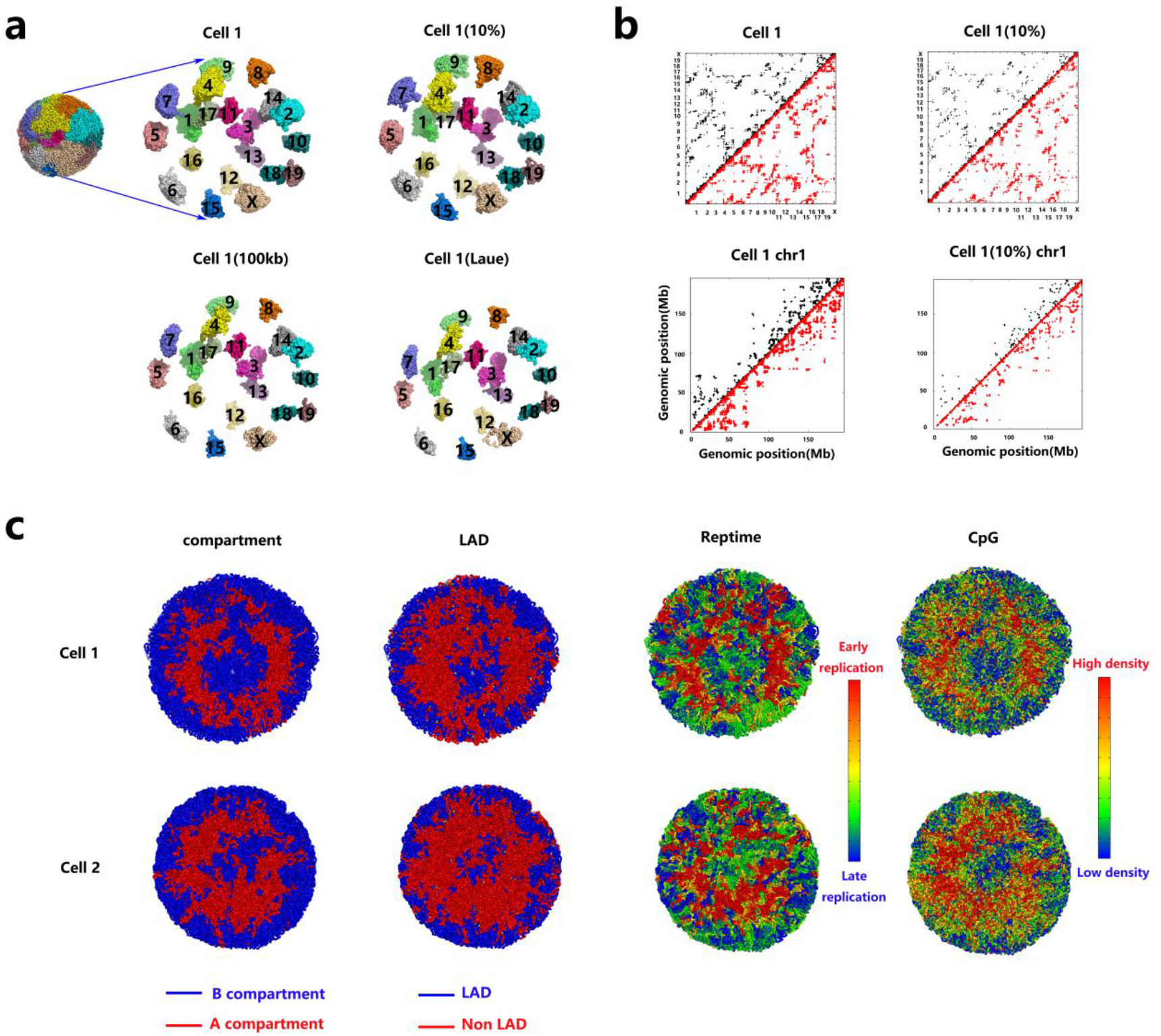
Validation of calculated structures. **a.** Expanded view of spatial territories for chromosomes in calculated structures of cell 1 constructed with different methods: 10kb resolution with all Hi-C data(upper left), 10kb resolution with 10% Hi-C data(upper right), 100kb resolution with all data(lower left) and 100kb resolution structures presented in Stevens’^29^ work(lower right). **b.** Comparison between experimental Hi-C contacts (upper triangle, in black) and reconstructed contacts (lower triangle, in red) among genome-wide loci for structures constructed from different datasets: cell 1 with all Hi-C data(upper left) and only 10% data separately(upper right). Same with two above matrices with enlarged view for chromosome 1. c. Cross view of 3D structures of intact genome of cell 1 and cell 2 with chromosome loci colored based on the corresponding features: loci belonging to compartment A(red) or B(blue); loci that are identified as LAD (blue) or not (red); the replication timing of each locus calculated based on ChIP-seq data; and CpG density of each locus.

### High resolution 3D genome organization under high resolution

The modeled chromatin 3D structures demonstrated similar folding conformations displayed by those presented in Stevens’ work^29^(Fig.1a, Fig.1c). For example, *Rabl* configuration, and discrete chromosome territories were observed in all cells. Constitutive lamina associated domains (LADs) are consistently organized at nuclear membranes or nucleolar periphery, while early replicated regions and CpG enriched loci are located between them. Besides, directed mapping of ChIP-seq data on 3D structures shows that chromosome segments marked by CTCF, H3K27me3, H3K27ac, H3K36me3, H3K4me1, H3K4me3, active enhancers and active genes are coherently enriched in the spatial regions with distance of 0.3-0.7 nuclear radius to the center(shown in Fig.2a, Extended Data Fig.3c). This suggests that euchromatin prefer to locate away from nucleolus and membranes where heterochromatin prefers to reside. In the calculated structures, each chromosome in cell nucleus is organized in a fractal globule manner and domains of varied genomic size can be observed in separation index matrix plot. These domains exhibit hierarchical structures of chromosome organization from hundreds of kilobases to tens of megabases (Fig.2b). Considering the calculations are performed based on Hi-C contact matrices with varied interaction patterns among genome-wide loci, such consistent distribution of chromosome segments among individual cells suggests that both experimental data and modeled structures successfully capture the coherent features of genome organization.

**Figure 2.**
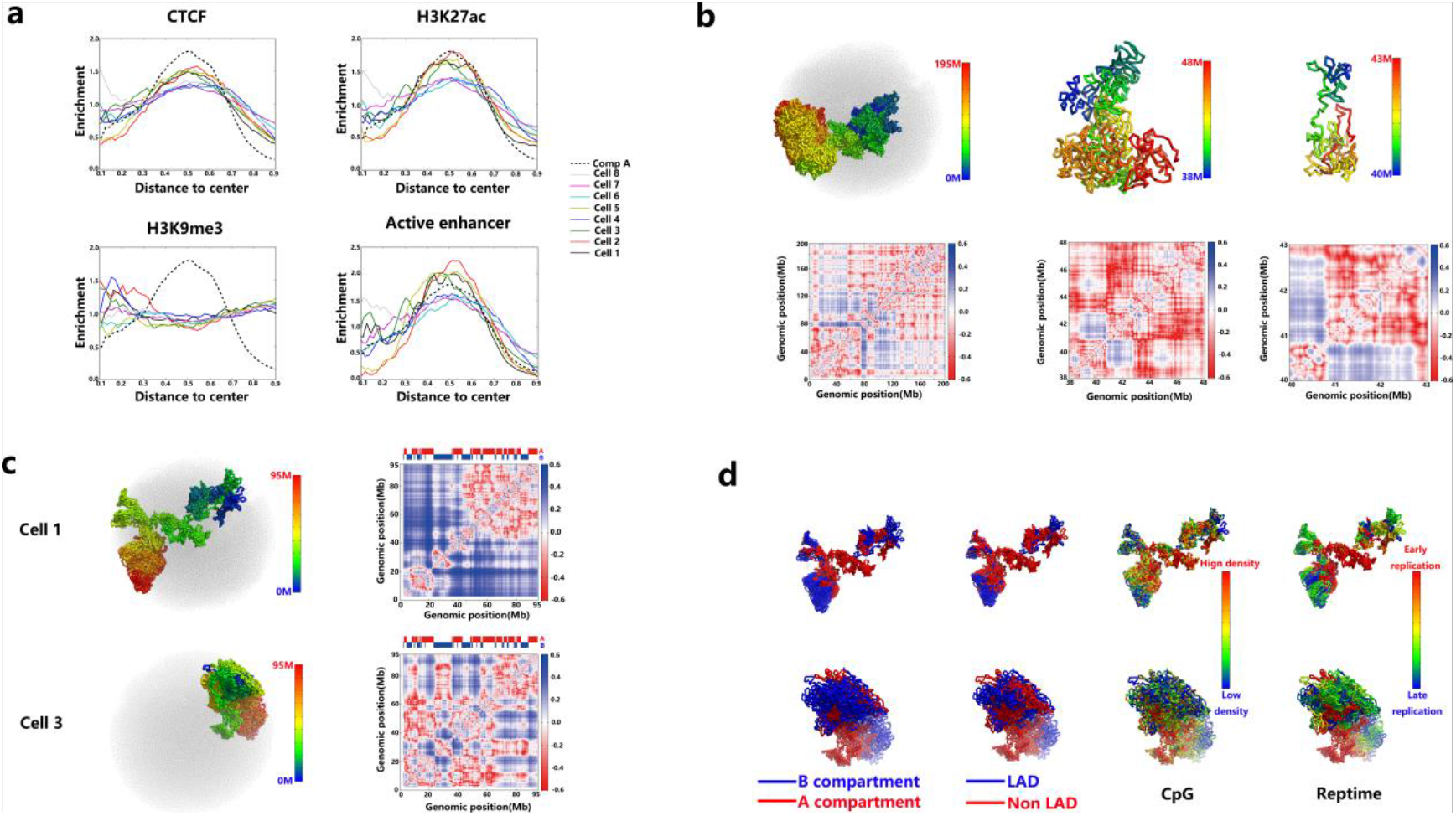
Consistent 3D organization of chromosome with cell dependent structures. **a.** Plot of enrichment (calculation presented in Supporting Information) of chromosome loci marked by CTCF (upper left), H3K27ac (upper right), H3K9me3 (lower left) and active enhancers (lower right)with respect to scaled radial distance to nuclear center in different cells. As comparison, enrichment of compartment A loci is also presented in dashed lines in each plot. **b.** Plot of 3D structures and the corresponding separation index matrices (bottom) for different chromosome regions in cell 1: intact chromosome 1 (left), 38M-48M in chromosome 1 (middle) and 40M-43M in chromosome 1(right). In 3D structure plot, chromosome loci are colored based on the genomic positions along the sequence. **c.** Same with B for chromosome 17 in calculated structures for cell 1 and cell 3, respectively. **d.** 3D projection of different chromosome features on varied structures of chromosome 17 from cell 1 and cell 3. The coloring of chromosome loci is performed with the same format shown in in Fig.1c.

### Varied 3D genome structures in individual cells

Consistent with substantially varied interaction patterns in Hi-C contact maps, chromosomes adopt alternative structures in organizing their consecutive segments in different cells, resulting in significant variation in 3D folding at different genomic scales. Such large structural variation is not due to the lack of contact constraints, since similar conformations are observed for structures calculated from the same Hi-C data. One example of such variation is that *chr17* adopts compact structure with polarized organization for A/B compartment in cell 3, while exhibits snake-like structure in cell 1 in a narrow but long territory confined by other chromosomes (shown in Fig.2c, Extended data Fig.2c). The varied organization of chromosome segments results in substantially different distance between intra-chromosome loci, which can be visualized in the plot of separation index matrices (shown in Fig.2c). These matrices show that regions of the same compartment type are not always aggregated in single chromosome, indicating the principles of genome folding derived from populated Hi-C may not always follow in single-cell level. These findings suggest that one chromosome can adopt different structure in the genome folding process, which to some extend contradict with the well accepted understanding about the determination of folded genome structure on the biological functions. Therefore, it is urgent to integrate these two scenes into a consistent picture that can unravel the underlying regulation mechanism of chromosome spatial organization in gene expression.

### Aggregation of chromosome regions with different features

As mentioned above, chromosome loci marked by different proteins, such as CTCF and H3K27me3, tend to spatially co-enriched with compartment A (Fig.1c, Extended Data Fig.3c), and show consistent aggregation in different cells (Fig.3a). Further analyses indicate that such aggregations of these loci are positively correlated with the increased genomic distance, regardless whether the marked proteins are same or not (Fig.3d, Extended Data Fig.4). This suggests the significance of proteins in regulating 3D genome structure. Strikingly, clustering analysis on loci marked by same protein shows that the aggregation are relatively randomly formed in different cells, without specific recognition on the genomic locations of loci in close spatial proximity (Fig.3b). Interestingly, H3K9me3 marked loci show relative weak aggregation with themselves but significant strong aggregation with those marked by other proteins (such as H3K27me3, H3K36me1, H3K4me1 and H3K4me3) (Fig.3a, Fig.3d). For loci marked by different proteins, coherent spatial aggregations are observed in different cells; it not only suggests the wide distribution of such aggregation in genome folding, but also further validates the reliability of our calculated structures. These consistent organizations suggest that the non-specific aggregation of protein marked loci may play important roles in maintaining coherent biological functions of individual cells of the same type.

**Figure 3.**
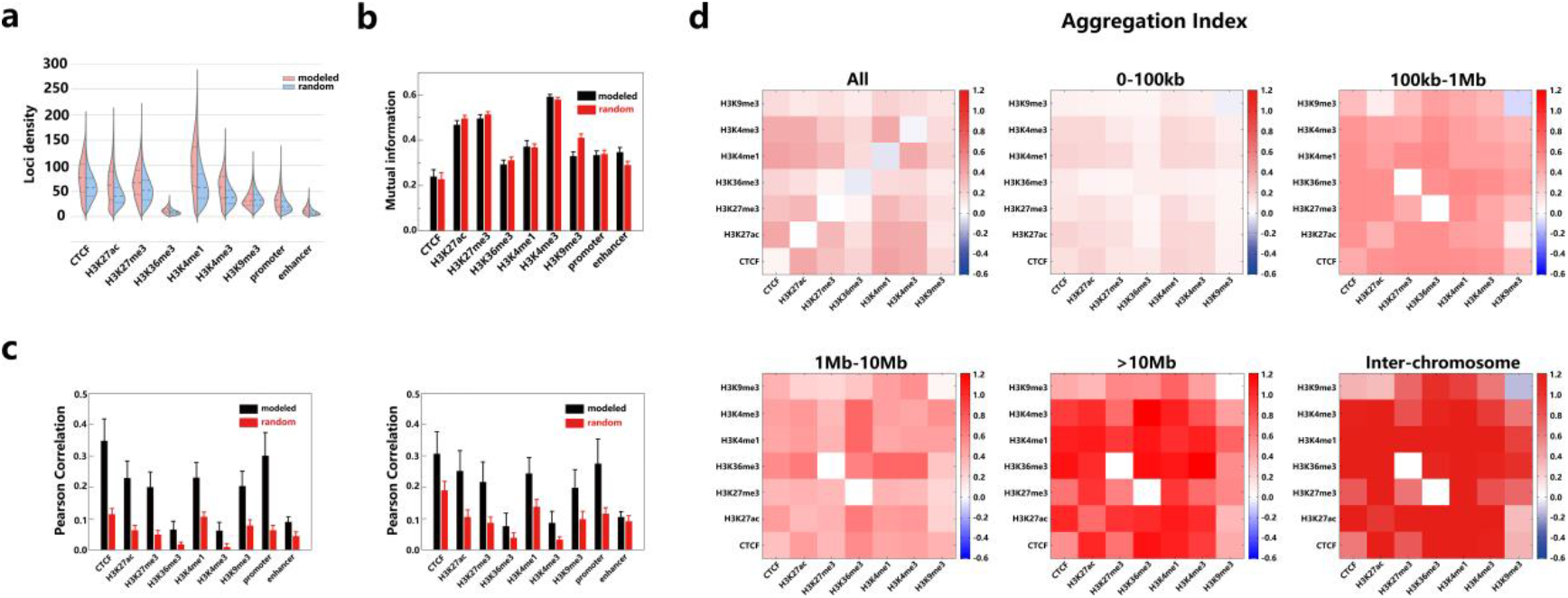
Non-specific aggregation of chromosome loci marked by proteins. **a.** Violin plot of the distribution of local density of chromosome loci marked by different proteins in cell 1. Relative high local density in modeled structure indicates these loci are aggregated in folded genome. **b.** Mutual information between spatial clusters formed by spatially aggregated loci marked by the same protein in different cells. As comparison, similar calculation is performed for random structures. Similar mutual information values suggest spatial clusters of aggregated proteins are randomly formed in modeled structures. **c.** Correlation between spatially averaged protein density (loci within 1Mb excluded in calculation) and sequentially averaged protein density (smoothed over 100kb) (left); Correlation between spatial averaged protein density (loci within 1Mb excluded in calculation) from different cells (right) **d.** Heatmap plot of aggregation degree matrix for chromosome loci marked by different proteins with respect to varied genomic distances. All calculations are performed based on modeled structures of all cells except panel (a).

In addition to the spatial aggregation of loci marked by different proteins, the average protein densities smoothed over loci in close spatial proximity show strong correlation with the average densities smoothed over genomic sequence (Extended Data Fig.5a). Similar results are obtained after excluding loci within 1Mb regions when selecting spatially close loci (Fig.3c), proving that such correlation is not caused by the coherence between spatial proximity and sequential proximity. Considering protein density is obtained based on ChIP-seq data of populated cells, such strong correlation suggests this property is robust among individual cells. Also, significant strong correlations of spatially averaged protein densities are observed between different cells (for CTCF average correlation value is 0.31, compared to 0.19 for random structures) (Fig.3c), suggesting that each locus is surrounded by chromosomes of similar biological features. The varied 3D organization of genome and consistent local biological environment of chromosome loci suggests that it is the biological features of the surrounded chromosome regions, instead of the exact conformation of folded chromosome, that maintain the similar biological functions in same type of cells. We therefore hypothesis that it is the non-specific aggregations between loci marked by different proteins may contribute to achieving such consistent biological environment.

### Association between active enhancers and active promoters with protein marked loci

The spatial association between enhancers and promoters, which has been validated by 3C experiments^30,31^, is thought to regulate the gene expression levels. In our modeled structures, an average of 2.57 active enhancers are located in close spatial proximity (within cutoff distance of 4 particle radii) from one active promoter, whereas only 0.04 contact pairs on average can be detected by current single cell Hi-C experiments. The spatial aggregation between active promoters and active enhancers is supported by experimentally observed structures such as transcription factories^32,33^, and active chromatin hub^3,34,35^, as well as recently proposed phase separation model^36–38^, which independently validate our observation that is from the computational perspective. Different from experimentally identified proximate pairs between active enhancers and promoters, which are mostly located within genomic distance of 1Mb, the spatial association in calculated structures can be formed with genomic distance up to hundreds of megabases (Fig.4a). Compared to random structures, the number of spatially associated active enhancer-promoter pairs with genomic distance greater than 1Mb (including inter-chromosome) has increased by 110%, suggesting remarkable promotion of such associations in genome folding (Fig.4a). For calculated cells, the associated pairs with genomic distance larger than 1Mb cover 43.9% active genes, suggesting long range enhancer-promoter association can also play an important role in regulating gene expression in cells. Moreover, the calculated structures show that the associated enhancer-promoter pairs vary substantially from cell to cell (Fig.4b) (an average of 0.05% common enhancer-promoter pairs with genomic distance larger than 1Mb are observed in two cells), which agrees with the features of super-enhancers^39,40^ that can regulate the expression of different genes in populated cells. The high percentages of active promoters that are non-specifically associated with active enhancers indicate varied folded genome structure can achieve similar gene activation profiles in individual cells.

**Figure 4.**
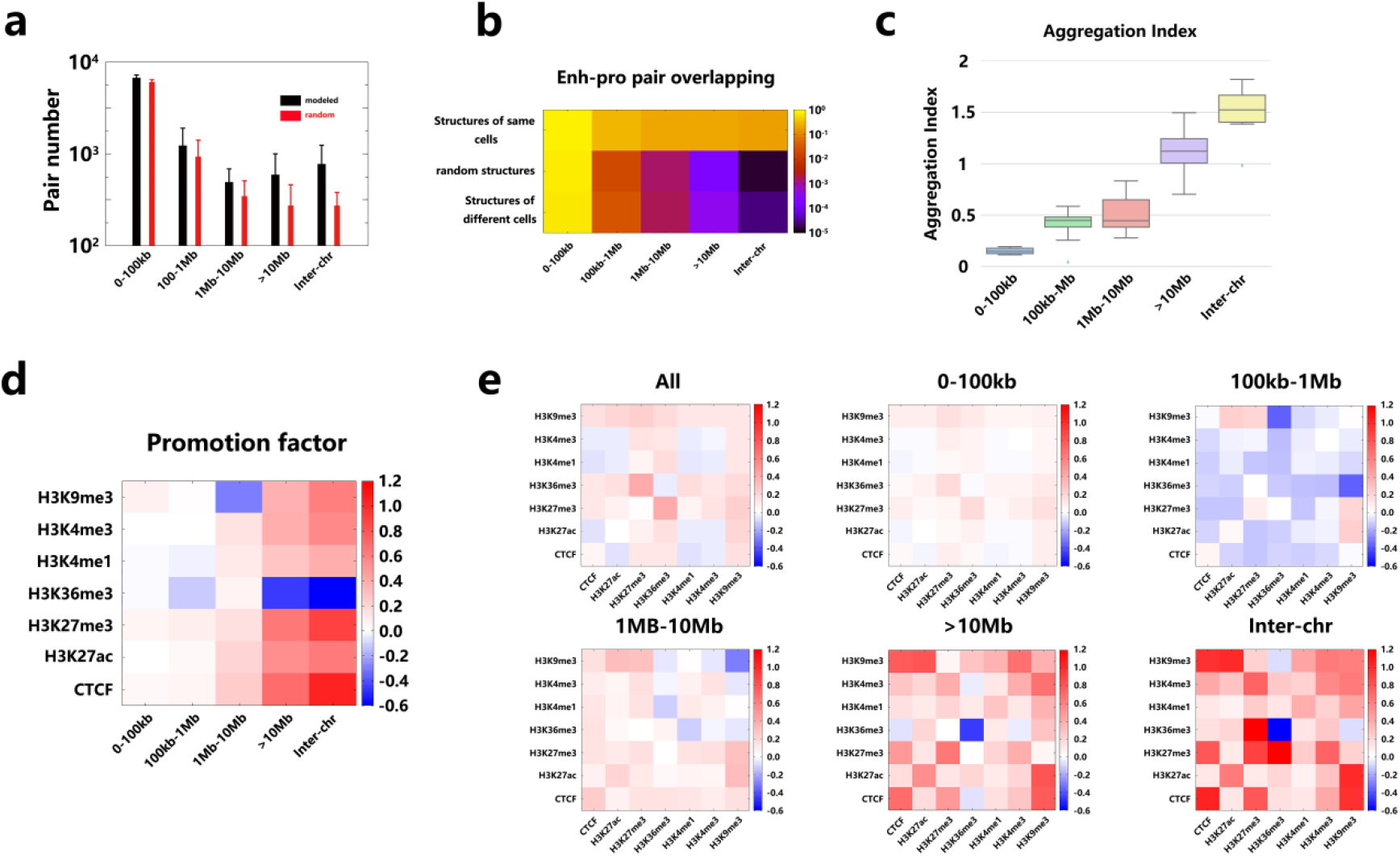
Spatial aggregation between active enhancers and promoters correlate with aggregation of chromosome loci marked by proteins. **a.** Comparison between aggregated active promoter-enhancer pair number for calculated structures and random structures with respect to different genomic distance. Calculation was performed on all cells. **b.** Heatmap plot of overlapping ratio between aggregated active promoter-enhancer pairs from different structures: structure replicas of the same cell (top row), random structures for different cells(median row) and modeled structures from different cells (bottom row). **c.** Box plot of aggregation degree between active promoters and active enhancers based on calculated structures for 8 cells with respect to different genomic distances. **d.** Heatmap plot of promotion factor of active enhancer-promoter association with respect to spatial aggregation of loci marked by the same proteins. **e.** Plot of promotion factor for chromosome loci marked by different proteins with respect to different genomic distances.

To explore the influence of aggregation of protein marked loci on the association between active enhancers and promoters, we defined a promotion factor that quantifies the improvement of enhancer-promoter association under the condition that two loci must be marked by the analyzed proteins. The result shows that the spatial aggregation between loci marked by proteins such as CTCF, H3K27ac, H3K27me3, H3K4me1 and H3K4me3 significantly promotes the active enhancer-promoter association and the promotion effect grows with the increase of genomic distance (Fig.4d, Fig.4e). Interestingly, H3K9me3 inhibits these associations in genomic distance 1Mb to 10Mb, while exhibit promotion effect for distance greater than 10Mb. The varied promotion patterns exhibit different roles of protein in regulating enhancer-promoter association. Similarly, aggregation of loci marked by different proteins show inhibition effect for distance in 100kb-1Mb region while promote active enhancer-promoter association for distance larger than 10Mb with different degrees (Fig.4e). These results suggest that the association between active enhancers and promoters are correlated with the aggregation network among loci marked by different proteins.

## Conclusion

In this work, we developed a new method to model experimental consistent high-resolution (10kb resolution) three-dimensional structures for intact genome based on single-cell Hi-C data of 100-kb resolution. The generated chromatin 3D structure results allow us to explore the relationship between chromosome folding and gene regulation at the single nucleus level. The modeling results demonstrated both cell-specific folding conformations of the genome and consistent organization of chromosome regions labeled by different features. Based on these structures, we discovered that chromosome loci marked by different proteins, including CTCF and modified histones are observed to randomly aggregate in nuclear space to ensure each locus is spatially surrounded by loci of similar biological features in individual cells. Moreover, such random aggregation of these loci is also correlated with regulating the association between active promoters and active enhancers at different genomic scales, suggesting that consistent spatial aggregation between chromosome loci marked by proteins in varied 3D genome structures plays an important role in maintaining similar gene expression levels of individual cells of the same type. Through this work, we would like to point out that theoretical analysis on 3D intact genome structures modeled from single-cell Hi-C data can effectively facilitate our investigation and understanding of chromosome organization, as well as the impact on biological functions of nucleus in single-cell level.

