## Supporting Information for "High resolution single-cell chromatin 3D modeling reveals coherent chromatin aggregation with varied structures in controlling genome function stability"

### Method

#### 1. Method to construct high resolution genome structures

3D genome structure calculation were conducted by applying molecular dynamics (MD) simulations to optimize the initial conformation of intact genome to satisfy the constraints of contact reads obtained in single cell Hi-C experiment. In the MD simulation, each chromosome was represented by one polymer chain with monomers denoting chromosome loci of equal genomic length. Software is available online https://github.com/TheMengLab/single_cell_hapliod with details described as below.

Initial filtering of isolated contact read: Similar with the filtering in Stevens'[1] work, Hi-C contact read with genomic distance larger than 2 Mb to any other reads is considered as isolated one in this work, which has relative high risk of sequence mapping errors. Therefore, these reads were removed in the whole calculated process to achieve more reliable Hi-C data in restraining geometric conformation of the whole genome.

Obtaining subset of Hi-C data (or removing certain percentage of contact reads from Hi-C data): For one cell, a subset of Hi-C data was obtained by randomly selection of contact reads among genome-wide segments with certain probability. In this process, the same probability was adopted for contact reads between intra-chromosome loci and inter-chromosome loci. The selected contact reads were treated as a new set of Hi-C data after random removal.

Merging genome wide contacts into contact frequencies between loci of same size: As mentioned above, polymer model was adopted for the presentation of single chromosome and each monomer (or particle) representing one chromosome segment of the same genomic size. Hi-C data after isolated read filtering were assigned to contact reads between pairs of monomers based on the genomic locations of the restriction fragment ends. Contact reads within the same monomer were ignored since they provide no information of polymer structural organization. The genomic size of each monomer varied with respect to the final resolution of modeled structure. For example, in the calculation of 10-kb resolution structure modeling, the structures were constructed with each monomer representing 1280 kb, 640 kb, 320 kb, 160 kb, 80 kb, 40 kb, 20 kb and 10 kb, separately. For the calculation of 1kb-resolution structure modeling, the genomic size were 1024 kb, 512 kb, 256 kb, 128 kb, 64 kb, 32 kb, 16 kb, 8 kb, 4 kb, 2 kb and 1 kb.

Void regions identification: In single-cell Hi-C data, some genome regions that no contact reads could be mapped to were considered as void regions in our calculation. These void regions were difficult to be modeled because of the absence of spatial constraints(except the constraints from sequential connectivity) with other genomic regions and were technically defined as those that have no contact reads with any regions in experimental data of all samples. Since the coordinates of these void regions may not be reliable, these regions were excluded from the further analysis of genome structure organization.

Force field between polymer particles: In this work, the units of mass, distance, time were all arbitrarily set as 1 (the particle radius of each monomer was set as 0.5). The force field of the system was designed to reproduce the constraints derived from experimental data as well as maintain the connectivity of polymer chain. For the connectivity maintaining, simple harmonic potential was adopted in our modeling with the formula

$$E_{bond} = k_{bond}\left(d - d_{eq}\right)^2$$

In this formula, d is the distance between two neighbor particles, $d_{eq}$ and $k_{bond}$ were predefined parameters with values set to 1 and 100, separately.
For the second term, we adopted different forms for two particles by considering whether contact reads C between them is zero or not. If C>0, it indicates two particles should be located at close spatially proximity and the energy function is designed as

$$E_1(d) = \begin{cases} \left(-0.667 * d - \frac{0.3}{4} ln\left(1 + e^{-4(d-2)}\right) + \frac{C}{2} ln\left(\frac{2+d}{2-d}\right)\right) * 10, & if\ d \leq 1.99 \\ \left(-0.667 * d - \frac{0.3}{4} ln\left(1 + e^{-4(d-2)}\right) + C(50d - \alpha)\right) * 10, & if\ d > 1.99 \end{cases}$$

In this formula, α is set as 96.5 to make the energy function to be continuous at d=1.99.

On the other hand, if C=0, it indicates two particle have higher probability to be separated, the energy function is designed as

$$E_2(d) = \begin{cases} \left(-2\ln d + 0.667 * d + \frac{0.3}{4} ln\left(1 + e^{-4(d-2)}\right)\right) * 10, & if\ d \leq 3.0 \\ \beta * 10, & if\ d > 3.0 \end{cases}$$

Similarly, β is set as 0.105 to make the energy function to be continuous at d=3.0.

Structure optimization: Starting from initial structure, the genome conformation was optimized by integrating Newtown's equation with pre-designed potential energy (described above) applied on all particles aimed at satisfying the constraint of experimental observation. In one integration step, force on each monomer was firstly obtained by differentiating these energy functions in 3 directions and these forces from different monomers were accumulated to calculate the total force on the monomer. After the force calculation, the maximum force value among all the monomers could be determined, which was noted as $F_{max}$. Then the time interval for numerical integration *dt* was calculated by $\mathrm{dt} = \sqrt{0.2/F_{max}}$. Such determination of *dt* can effectively improve the efficiency of structural optimization. Finally the conformation of the polymer was updated by integrating Newtown's equation with calculated forces and time interval. The velocity of each particle was randomly generated in each step. Each optimized conformation was obtained after 10000 integration steps from initial structure.

Hierarchical scale calculation to obtain high-resolution structures: The genome structure was hierarchically constructed from low resolution to high resolution (the genomic size of each monomer in one structure changes from 1280 kb to 640 kb, 320 kb, 160 kb, 80 kb, 40 kb 20 kb and eventually 10 kb). The initial structure of 1280 kb resolution was randomly generated, which was optimized following the procedure of integrating Newtown's equation. The optimized 1280kb resolution structure was then used to generate initial structure of 640 kb resolution, which was also optimized following the same procedure. Using this method, we iteratively obtained optimized structures of different resolutions until the 10-kb resolution structure. To remove bias introduce by the initial structures, 20 10-kb structure replicas were generated starting from different initial structures at 1280-kb resolution for structural validation and statistics. Calculation of structure of 1kb resolution is performed in similar protocol from initial resolution of 1024 kb.

#### 2. Contact matrix reconstruction

Reconstruction of contact matrix was performed based on the optimized structures by checking whether two monomers were close enough or not. In our calculation, binary contact value between pair of loci was adopted by considering the spatial distance between them: contact value of 1 for spatial distance smaller than $d_{cutoff}$=2 (4 times of particle radii) and 0 for distance larger than $d_{cutoff}$. Loci pairs that were separated with distance larger than $d_{cutoff}$ are considered as violated pairs if experimental contact reads existed between them. For each single cell, the calculation of contact matrix reconstruction is performed on one structure replica that was randomly selected.

#### 3. Source list of experimental data and pre-calculation

The experimental data used in our analysis is taken from previously published work, as elaborated below:

| Data type | Accession number | Reference |
|---|---|---|
| CTCF ChIP-seq (haploid) | GSE80280 | Stevens, T. J. *et al.* 3D structures of individual mammalian genomes studied by single-cell Hi-C. *Nature* **544**, 59-+, doi:10.1038/nature21429 (2017). |
| Active enhancer/promoter | GSE29184 | Shen, Y. *et al.* A map of the cis-regulatory sequences in the mouse genome. *Nature* **488**, 116-120, doi:10.1038/nature11243 (2012). |
| H3K27ac | GSE56098 | Buecker, C. *et al.* Reorganization of Enhancer Patterns in Transition from Naive to Primed Pluripotency. *Cell Stem Cell* **14**, 838-853 (2014). |
| H3K27me3 ChIP-seq (haploid) | GSE80280 | Stevens, T. J. *et al.* 3D structures of individual mammalian genomes studied by single-cell Hi-C. *Nature* **544**, 59-+, |

| | | |
|---|---|---|
| | | doi:10.1038/nature21429 (2017). |
| H3K36me3 | GSE80280 | Stevens, T. J. *et al.* 3D structures of individual mammalian genomes studied by single-cell Hi-C. *Nature* **544**, 59-+, doi:10.1038/nature21429 (2017). |
| H3K4me1 | GSE56098 | Buecker, C. *et al.* Reorganization of Enhancer Patterns in Transition from Naive to Primed Pluripotency. *Cell Stem Cell* **14**, 838-853 (2014). |
| H3K4me3 ChIP-seq (haploid) | GSE80280 | Stevens, T. J. *et al.* 3D structures of individual mammalian genomes studied by single-cell Hi-C. *Nature* **544**, 59-+, doi:10.1038/nature21429 (2017). |
| H3K9me3 ChIP-seq (haploid) | GSE80280 | Stevens, T. J. *et al.* 3D structures of individual mammalian genomes studied by single-cell Hi-C. *Nature* **544**, 59-+, doi:10.1038/nature21429 (2017). |
| Constitutive Lamina Associated Domain | GSE17051 | Peric-Hupkes, D. *et al.* Molecular Maps of the Reorganization of Genome-Nuclear Lamina Interactions during Differentiation. *Mol Cell* **38**, 603-613, doi:10.1016/j.molcel.2010.03.016 (2010). |
| Replication Timing | E-MTAB-3506 | Kolesnikov, N. *et al.* ArrayExpress update-simplifying data submissions. *Nucleic Acids Res* **43**, D1113-D1116, doi:10.1093/nar/gku1057 (2015). |

For the ChIP-seq data that provide the information of protein enrichment at different genomic regions was first mapped to the genome wide loci presented as monomers in calculated structures. The mapped density on each locus is calculated as the averaged signal(weighted by the genomic length of detected DNA sequence) of sequence located in the corresponding genomic region. After the mapping, the loci with non-zero protein density were noted as protein marked ones.

#### 4. Identification of A/B compartment

The identification of chromosome compartment was calculated following a similar algorithm described in previous work[2], in which correlation matrix between normalized contact frequency matrix was calculated and PCA algorithm was adopted to dividing one chromosome into two compartments without assignment for compartment types (in total 40 compartments for 20 chromosomes). Then inter-chromosomal contact frequency matrix between these 40 compartments was calculated, which were further scaled by the genomic length of each compartment to eliminate the bias introduced by varied loci number in each region. Finally, regions with higher normalized inter-chromosome contact frequency value were assigned as A type, and the other regions are assigned as B compartment type. The Hi-C matrix used is taken from previous work[1], and a resolution of 200 kb was adopted in identifying compartments.

5. **Enrichment of chromosome loci marked by protein.**

The enrichment of chromosome loci marked by protein is defined as the number of chromosome loci in specified spatial regions scaled by the corresponding loci number in the same regions from random structures. Enrichment values that are larger/smaller than 1 separately suggest the protein marked loci is relative enriched/depleted in that spatial region along the radial direction.

6. **Spatial density of protein marked loci**

The spatial density of one protein marked locus was defined as the number of loci located in spatial proximity, as well as marked by the same protein. In calculation, the cutoff distance for spatial proximity was set as 10 particle radii. For one individual cell, the spatial densities of chromosome loci were averaged from the density values calculated from different structure replicas modeled from the same experimental data.

7. **Calculation of scaled radial distance to nuclear center**

The varied structures calculated for different cells suggested that the constructed genomes have the risk of varied nuclear radii, which would introduce some inconsistency when comparing the distribution of protein marked loci along the radial direction to nuclear center. Therefore, we rescaled the radial distance between one locus and nuclear center by the nuclear radius (defined as the maximum value of radial distance among all loci) to generate the scaled radial distance. After the scaling, radial positions of chromosome loci from different cells are comparable in the analyzing the organization of genome folding.

8. **Calculation of separation index matrix**

The separation index between two loci was used to describe the relative tendency of two loci to be spatially separated when compared to genome wide loci pairs separated by regions of the same genomic length. Mathematically, the separation index S was defined as $S = \frac{\log(d/d_{ave})}{log10}$, in which $d$ represents the spatial distance between two loci and $d_{ave}$ denotes the genome-wide averaged distance for loci pairs separated by the same genomic distance. The separation index matrix was constructed with each entry representing the separation index between two corresponding loci. Heatmap plot of separation index matrix provided direct visualization of chromosome folding at chromosomal scale.

9. **Calculation of aggregation degree between protein marked loci**

The aggregation degree was used to describe the extent of aggregation level between loci marked by specified features, which was defined as the logarithm ratio between the number of spatially aggregated loci pair in modeled 3D structures and the corresponding aggregated pair number in random structures. The formula of aggregation degree is expressed as $D_{agg} = \frac{log\left(N_{model}/N_{random}\right)}{log2}$, in which $N_{model}$ and $N_{random}$ separately represented the number of aggregated loci pairs in calculated 3D structures and random structures. In the calculation, two loci were considered to be aggregated if the corresponding spatial distance was less than 4 particle radii. The more positive Dagg is, the stronger extent of aggregation of loci marked by certain protein. In addition, we separately calculated the aggregation degree for loci pairs separated by different genomic distances to explore the organization of protein marked loci at different scales. In this work, 5 classes of genomic distances were adopted: 0-100 kb, 100 kb-1 Mb, 1 Mb-10 Mb, >10 Mb (intra-chromosomal) and inter-chromosome.

10. **Generation of random structures**

The random structures were obtained by shifting the genomic positions of each monomer in calculated 3D structure with a circular permutation manner along chromosome sequence, while keeping the coordinates of monomers unchanged. The shifted distance along genome sequence was set to arbitrary large values, such as 500Mb, 1Gb, 1.5Gb and 2Gb.

11. **Identification of spatial clusters of protein marked loci**

The spatial clustering of chromosome loci marked by the same protein was performed with the density peak based algorithm[3]. In this clustering algorithm, two parameters needed to be defined: spatial density of protein marked loci and cutoff criteria for determining centers of clusters. In this work, the spatial density of protein marked loci $\rho$ was calculated with definition described above and chromosome loci were defined as cluster centers if its spatial density $\rho$ and the nearest distance to the locus of higher spatial density $\mathrm{d}$ satisfies $\rho * \mathrm{d} > \frac{N_{protein}}{5}$, in which $N_{protein}$ represent the total number of loci marked by the protein. The distance $\mathrm{d}$ here is calculated in length unit(1 particle radius=0.5 length unit). Similar calculations were taken for identifying spatial clusters in random structures.

12. **Mutual information between different cluster features**

Mutual information is a statistical parameter to quantify the similarity between two sets of assignments for clusters on the same data. In this work, mutual information was adopted to measure the similarity between assigned clusters for chromosome loci marked by the same protein in structures X and structure Y. In structure X, suppose the protein marked loci were partitioned into $N_X$ clusters with $p(x)$ representing the percentage of loci assigned to the xth out of $N_X$ clusters. Similarly for structure Y, the corresponding cluster number is $N_Y$ and $p(y)$ noting the percentage of loci assigned to the yth out of $N_Y$ clusters. In addition, based on the cluster assignments for two structures, $p(x, y)$ was defined as the percentage of marked loci that were assigned to the xth cluster in the structure X while were assigned to the yth cluster in structure Y. Then the mutual information between two clustering results was calculated with the formula $\mathrm{M}(X,Y)=\sum_{x=1}^{N_X}\sum_{y=1}^{N_Y}p(x,y)log\frac{p(x)p(y)}{p(x,y)}$. In our work, the mutual information between structure X and structure Y was further normalized by dividing the square root of the product between $\mathrm{M}(X,X)$ and $\mathrm{M}(Y,Y)$. The larger the normalized mutual information is, the more similar two sets of clusters are. For example, if two clustering are identical, the normalized mutual information will be 1 after the calculation.

**13. Spatial averaged feature intensity(or protein density)**

The feature intensity on each locus is numerically taken from marked protein density directly mapped from ChIP-seq data. The spatially averaged feature intensity was defined as the averaged protein density of loci located within spatial proximity of 10 particle radii. Different from locus density calculation described above, all chromosome loci satisfying the geometric constrains were used for calculation, regardless whether these loci were marked by protein or not. In addition, we also calculated the sequential-neighbor-excluded spatially averaged feature intensity by excluding the contribution of chromosome loci located with genomic distance less than 1Mb.

**14. Promotion factor calculation**

Promotion factor was defined to describe the improvement of active enhancer-promoter aggregation degree under the condition of spatial aggregation of protein marked loci, which was expressed with the formula $F_{Pro}=\frac{log\left(\frac{N_{p-e,protein}}{N_{protein}}\Big/\frac{N_{p-e,protein,ran}}{N_{protein,ran}}\right)}{log2}$, in which $N_{p-e,protein}$ represented the number of aggregated loci pairs that are marked by promoters and enhancers, as well as proteins. $N_{protein}$ represented the total number of aggregated loci pairs marked by the proteins. $N_{p-e,protein,ran}$ and $N_{protein,ran}$ denoted similar meaning but for random structures instead of modeled structures. The distance cutoff for the spatial aggregation was also set to be 4 particle radii(same value with the cutoff for Hi-C contact). Similarly with aggregation degree calculation, promotion factor calculation could also be conducted for loci pairs separated by different genomic distances to explore the influence of protein aggregation on the association between active enhancers and promoters.

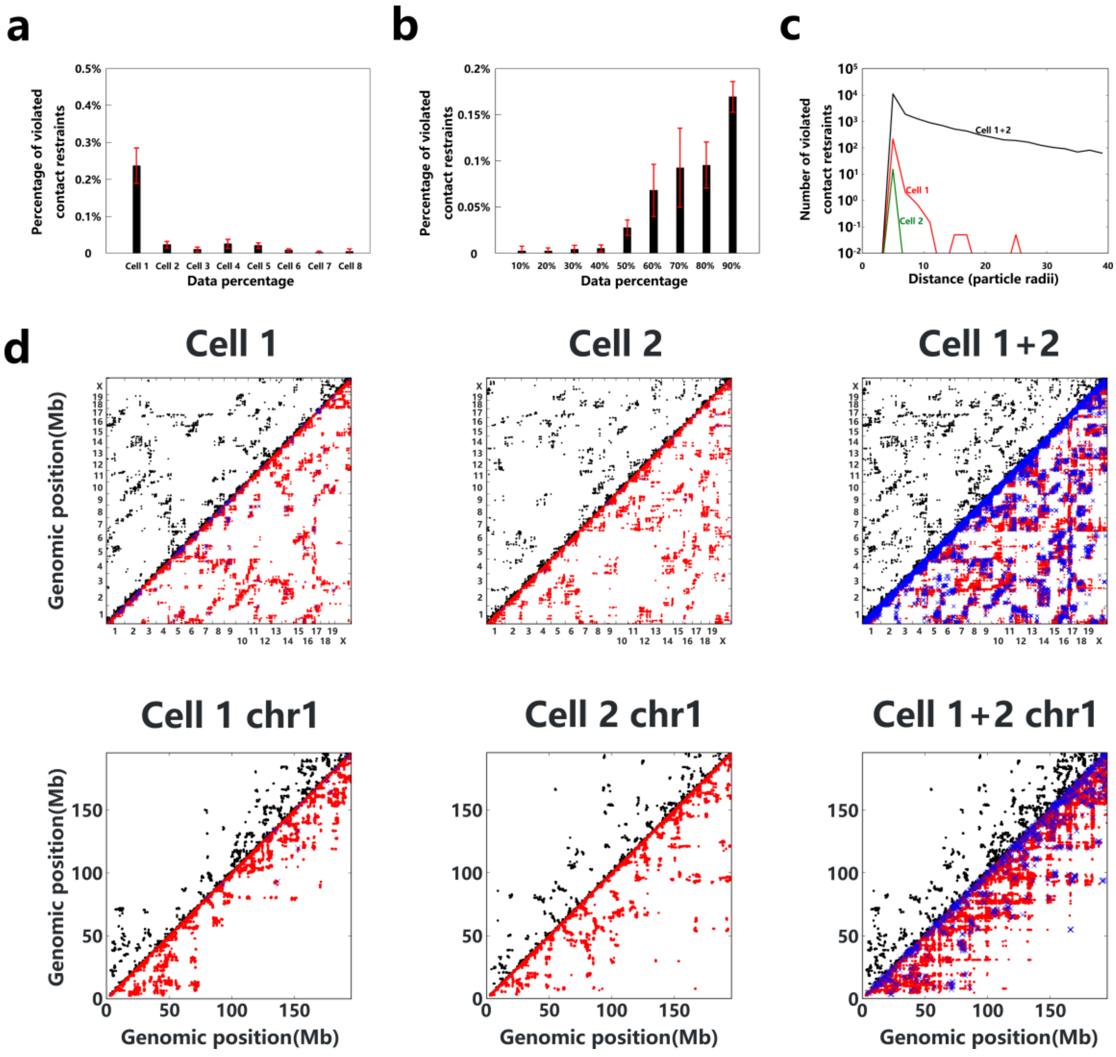


**Extended Data Figure 1 | Validation of modeling method in constructing 3D structures following Hi-C contact derived restraints. a,** Plot of the percentage Hi-C contact derived restraints that are not reproduced in calculated structures for different cells. **b,** Same with (a) for structures calculated based on dataset after randomly selecting subset of HiC contact reads in cell 1. The low percentage of violated contact derived restraints suggests the high quality of our method in constructing 3D genome structures for individual cells. **c,** The number of violated contact restraints in each structure for different Hi-C dataset: cell 1(red), cell 2(green) and combined Hi-C data of both cells (black). The result shows calculated structures from combined dataset have significant higher (by 1-2 magnitude) pair numbers as well as larger degrees in violating contact derived restraints. **d,** Comparison between experimental Hi-C contacts (upper triangle, black dot), reconstructed contacts (lower triangle, red dot) and violated loci pairs (lower triangle, blue cross) of constructed structures for cell 1, cell 2 and combined HiC dataset of two cells. Such result suggests that the loci pairs violating Hi-C contact derived restraints are more frequently located at the sequentially close regions in constructed structures. Also, the number of violated loci pairs increase dramatically after merging contact reads of two cells for 3D structure construction.

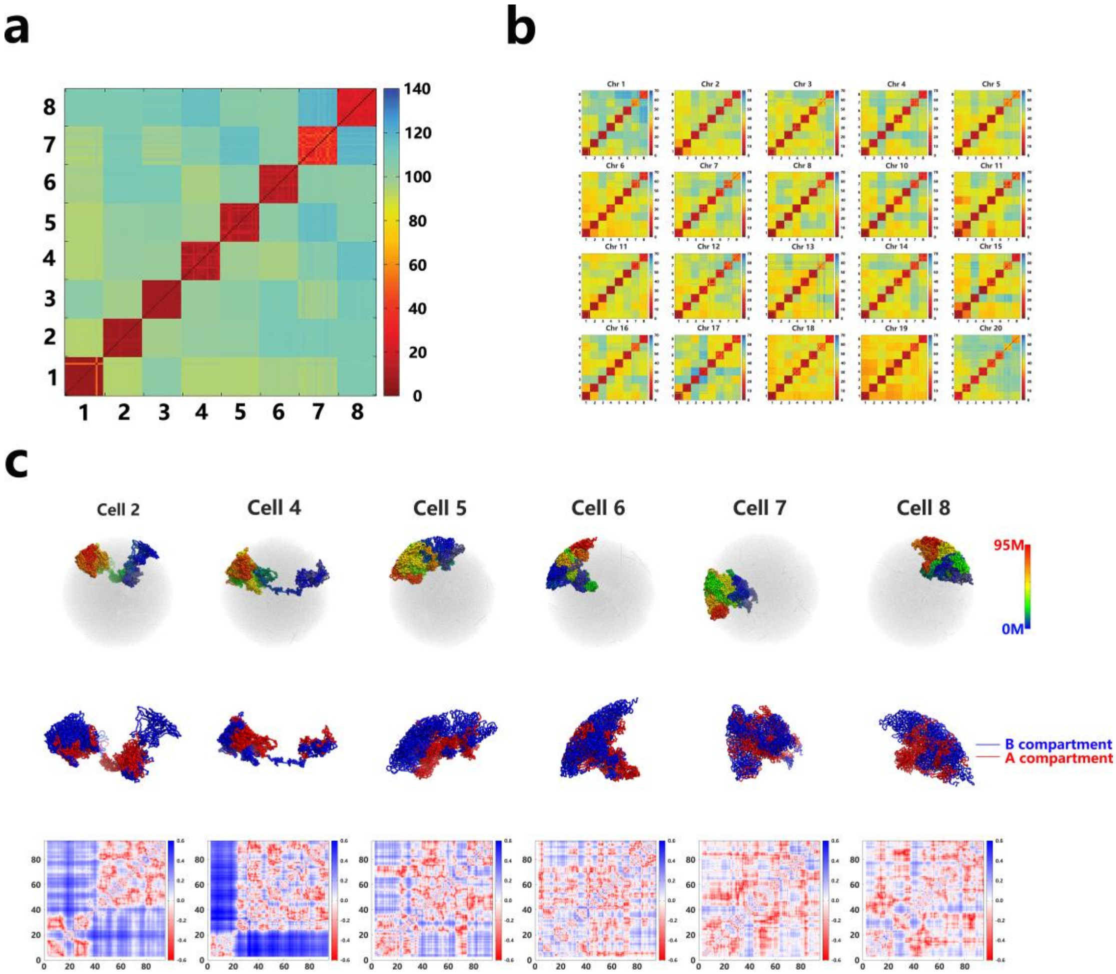


**Extended Data Figure 2 | Variation in modeled genome structures of individual cells. a,** Heatmap plot of root mean square deviation (r.m.s.d.) between intact genomes of all modeled structure replicas from 8 cells (in the unit of particle radii). **b,** Same with (a) for individual chromosome. **c,** Plot of 3D structures (top), compartment A/B (middle) and separation matrix (bottom) of chromosome 17 in different cells. These results suggest each chromosome adopt varied structures in genome folding of individual cells.

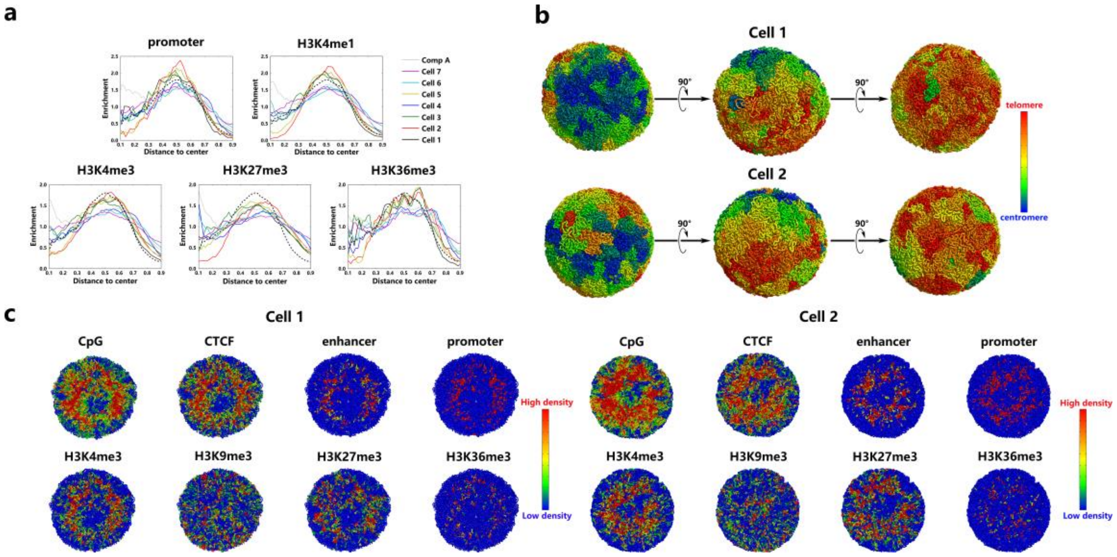


**Extended Data Figure 3 | Consistent organization of chromosome segments in individual cells. a,** Plot of enrichment of chromosome loci marked by protein with the same format presented in Figure 2(a) in main text. **b,** Consistent Rabl configurations are observed for cell 1 and cell 2. **c,** Cross view of 3D structures of intact genome of cell 1 and cell 2 with chromosome regions colored based on the averaged intensity of different features(smoothed over 100kb). Except H3K9me3 which exhibits relative random distribution, most features show enrichment at the spatial region between nucleolus and nucleolar peripheral region and such enrichment are conserved among individual cells.

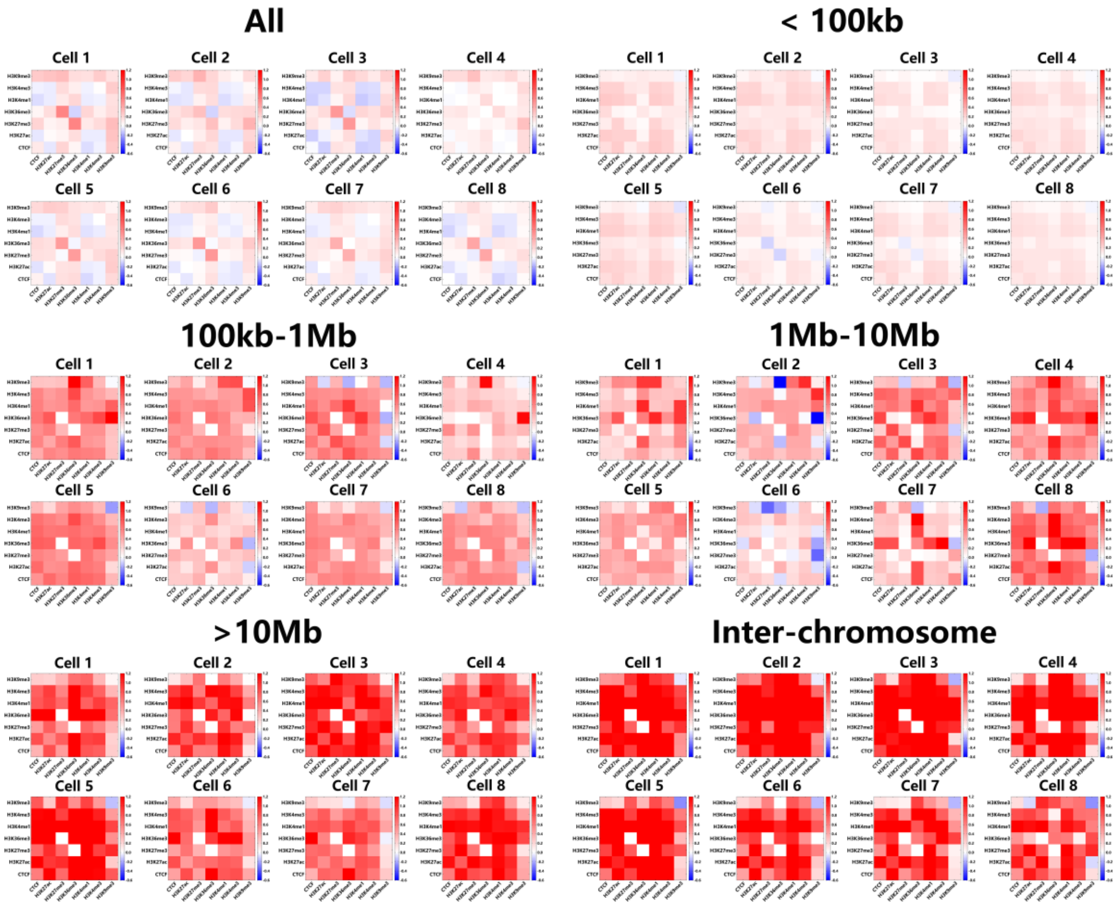


**Extended Data Figure 4 | Aggregation degrees between chromosome loci marked by different proteins in individual cells.** The aggregation degree increases with the increasing of genomic distance between loci marked by analyzed proteins, regardless the proteins are identical or not. The patterns of aggregation degree between different proteins are conserved among individual cells, suggesting the existence of principles for protein aggregation in controlling the folding of genomes.

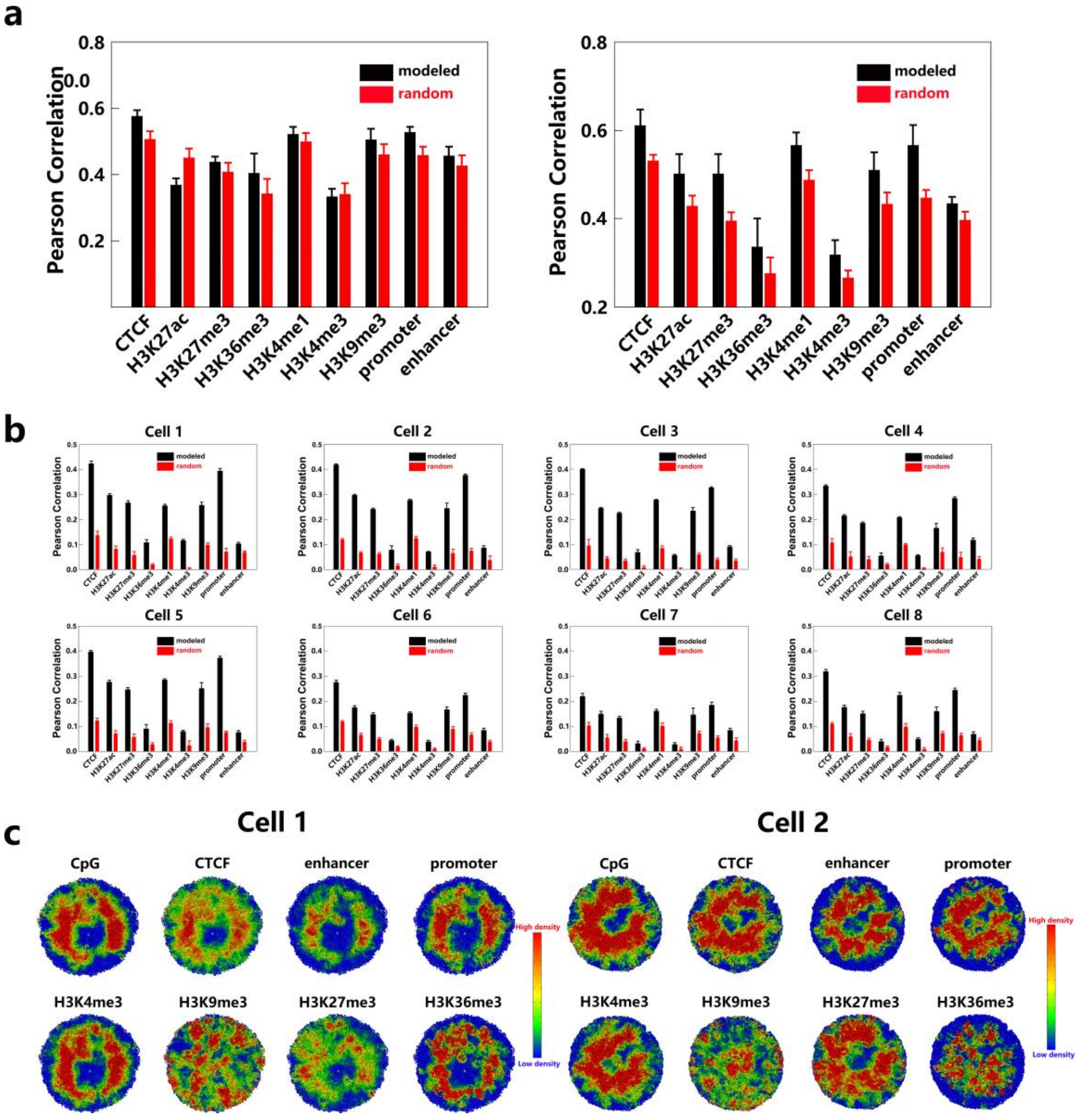


**Extended Data Figure 5 | Consistence chromosome averaged feature intensity separately smoothed over spatial regions and genomic sequence. a,** Correlation between protein density of spatial average (without sequential exclusion) and sequential average (smoothed over 100kb) (left); Correlation between spatial averaged protein density (without sequential exclusion) from different cells (right). **b,** Correlation between protein density of spatial average (loci within 1Mb excluded in calculation) and sequential average (smoothed over 100kb) for individual cells. **c,** Cross view of 3D projection of spatially averaged feature intensities (excluding sequentially proximity of 1 Mb) on calculated structures for cell 1 and cell 2. Compared to the projection for sequentially averaged feature intensities shown in Extended Data Figure 3(c), spatial averaged intensities also suggest the existence of nuclear central regions and nucleolar periphery that are depleted of marked proteins.

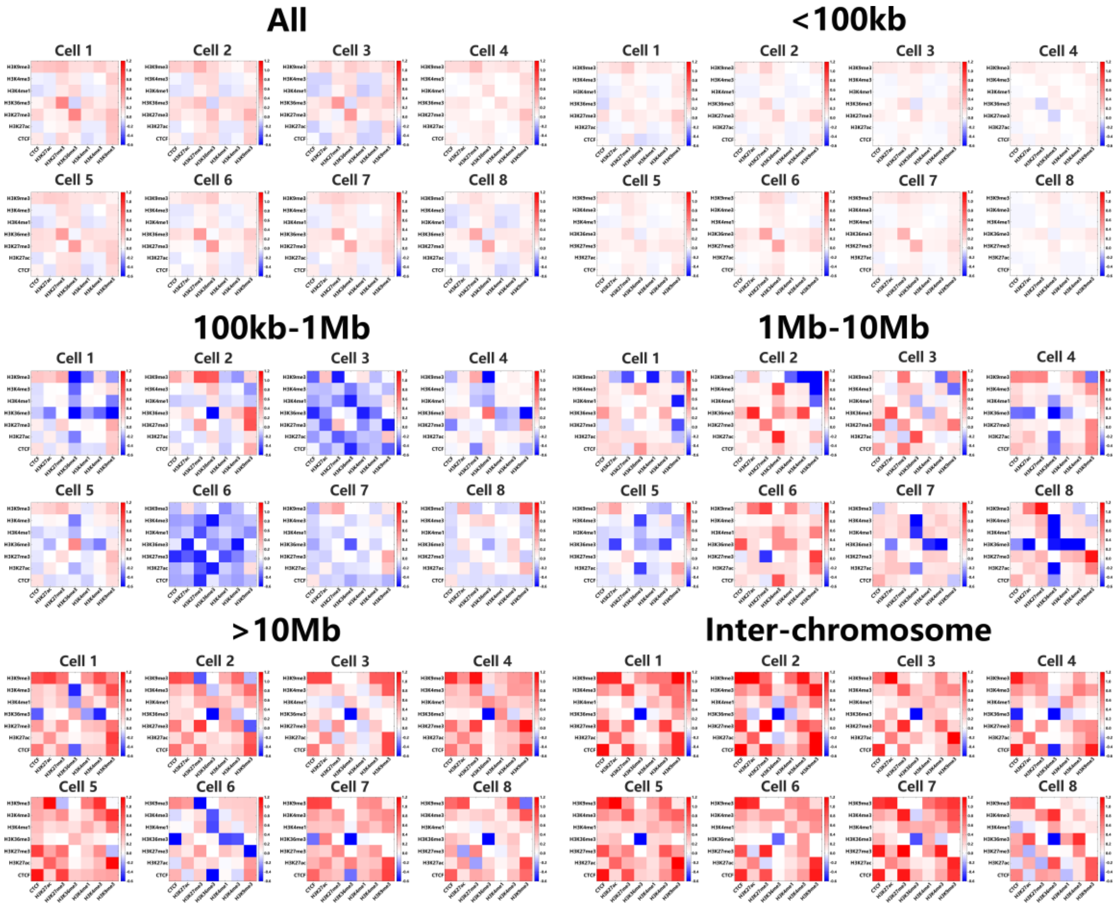


**Extended Data Figure 6 | Heatmap plots of promotion factor for the aggregation of active enhancers and promoters under the condition of spatial aggregation of different proteins in individual cells.** The heatmap plots show protein binding on DNA sequence can inhibit the aggregation of active enhancers and promoters for small genomic distance(for example, 100kb-1Mb), while promote such aggregations for sequentially far separated regions(for example, >10Mb or inter-chromosome).


1 Stevens, T. J. *et al.* 3D structures of individual mammalian genomes studied by single-cell Hi-C. *Nature* **544**, 59-+, doi:10.1038/nature21429 (2017).

2 Lieberman-Aiden, E. *et al.* Comprehensive Mapping of Long-Range Interactions Reveals Folding Principles of the Human Genome. *Science* **326**, 289-293, doi:10.1126/science.1181369 (2009).

3 Rodriguez, A. & Laio, A. Clustering by fast search and find of density peaks. *Science* **344**, 1492-1496, doi:10.1126/science.1242072 (2014).
